# TMEM41B Contributes to Atherosclerosis by Promoting Lipid Synthesis in Vascular Smooth Muscle Cells via Fatty Acid Synthase Stabilization

**DOI:** 10.1101/2025.03.24.644797

**Authors:** Gui-Yan Peng, Li-Tai Wei, Jun-Jie Ning, Ye-Xiang Jing, Hao-Long Luo, Zi-Lun Liu, Run-Chen Wei, Guang-Qi Chang, Mian Wang

## Abstract

Foam cell formation has traditionally been attributed to macrophages; however, emerging evidence highlights vascular smooth muscle cells (VSMCs) as another significant contributor. Here, we found TMEM41B is significantly upregulated in VSMCs of both human atherosclerotic (AS) lesions and murine models. VSMCs specific silencing TMEM41B expression in apolipoprotein E–deficient (ApoE−/−) mice markedly reduced plaque size and macrophage infiltration. Overexpressing TMEM41B in cultured VSMCs alters intracellular lipid profiles through stabilizing fatty acid synthase (FASN), a crucial enzyme in fatty acids synthesis, via inhibiting its ubiquitination and degradation. The TMEM41B/FASN axis drives lipid synthesis, promotes intracellular lipid storage, and facilitates the release of pro-inflammatory cytokines. Further, herpes simplex virus (HSV) infection amplified TMEM41B expression via OCT-1-mediated transcriptional activation, linking viral infection to lipid metabolic reprogramming in AS. These findings redefine the paradigm of VSMC-derived foam cell formation and suggest that targeting the TMEM41B/FASN axis could offer a transformative therapeutic strategy for AS, particularly in patients with HSV infection.

## 1 Introduction

Atherosclerosis (AS) is a major contributor to cardiovascular disease, and there is a steady rise of AS-related complications, with millions of individuals at risk for severe outcomes such as ischemic heart disease, stroke and lower limb ischemia[1–4]. Foam cell formation is a characteristic pathological feature of AS. The accumulation of foam cells secrete pro-inflammatory factors[5], along with subsequent apoptosis and necrosis, contributes to the formation of a necrotic core in unstable plaques, thereby increasing the risk of plaque rupture and thrombosis[6, 7]. Targeting foam cells formation may offer a promising strategy for the treatment and prevention of AS. Macrophages from bone marrow monocytes enter the vessel wall and absorb excess cholesterol[8], which have been considered the primary source of foam cells traditionally[9, 10]. However, recent evidence suggests that in atherosclerotic plaques, more than 50% of foam cells originate from vascular smooth muscle cells (VSMCs) rather than circulating monocytes[11, 12]. Despite this, the mechanism underlying the contribution of lipid-laden VSMCs to atherosclerosis remains largely unclear. Current studies on foam cell formation primarily focus on mechanisms such as scavenger receptor-mediated lipid uptake, lipid droplet degradation, and reverse cholesterol transport[9, 12–17]. Under pathological settings, VSMCs undergoes a phenotypic switch that can promote extracellular matrix synthesis[18, 19]. However, the role of intracellular lipid synthesis in VSMCs during these processes has been less explored.

As a multifunctional protein localized in the endoplasmic reticulum (ER), TMEM41B has been implicated in maintaining lipid homeostasis, as well as facilitating autophagosome formation and lipid droplet dynamics during the physiological state[20–23]. TMEM41B regulates these processes by controlling phospholipid distribution and mobilization to maintain cellular balance[20]. In addition to its role in lipid metabolism, TMEM41B is critical for viral replication, enabling ER membrane remodeling to support viruses like flaviviruses and coronaviruses[21, 24–26]. Emerging evidence suggests that pathological conditions, such as viral infections, may disrupt TMEM41B’s physiological function, transforming it from a homeostatic regulator to a driver of pathological lipid accumulation[27]. This dual functionality reflects the context-dependent complexity of TMEM41B in cellular regulation, although its contribution to pathological processes in atherosclerosis remains poorly understood. Understanding how TMEM41B contributes to lipid accumulation in VSMCs and AS progression could uncover new therapeutic targets.

In this study, we identified that TMEM41B is upregulated in VSMCs within atherosclerotic plaques. TMEM41B knockdown significantly alleviated AS progression in vivo. Mechanistically, TMEM41B promotes VSMC-derived foam cell formation through its direct interaction with fatty acid synthase (FASN), stabilizing FASN and enhancing fatty acid synthesis pathways. Furthermore, HSV infection activates the transcription factor OCT-1, upregulating TMEM41B expression and driving foam cell formation, thereby linking viral infection to metabolic reprogramming in AS. Collectively, this work suggests that the TMEM41B/FASN axis is essential for VSMC lipid accumulation and foam cell formation, providing a potential therapeutic target in AS.

## 2 Methods

### 2.1 Sample Collection

The study was approved by the Research Ethics Committee of the First Affiliated Hospital of Sun Yat-sen University, and informed consent was obtained from all participants. Detailed patient information is provided in Supplementary Table S1. Arterial samples were collected from six patients with severe lower extremity arterial disease who underwent amputation surgery, alongside six normal arterial samples obtained from healthy organ donors.

### 2.2 Animal

To minimize the effects of estrogen on atherosclerosis, only male mice were used in this study. Six-to eight-week-old male C57BL/6J and apolipoprotein E-deficient (ApoE^−/−^) mice were obtained from the Animal Center of the First Affiliated Hospital of Sun Yat-sen University and Zhuhai Best Bio-technology Co. Mice were housed under specific pathogen-free conditions with a temperature of 24 ± 2 °C, relative humidity of 55%-65%, and a 12-hour light/dark cycle. Food and water were provided ad libitum. Atherosclerosis was induced by administering a Western diet containing 45% fat (kcal/100 g) and 0.2% cholesterol (TROPHIC) for the specified durations. The animal studies were authorized by the animal experimental ethics committee of the First Affiliated Hospital of Sun Yat-sen University.

Adeno-associated virus 9 (AAV9) vectors expressing short hairpin RNA (shRNA) targeting TMEM41B (AAV-shTMEM41B) and scrambled control sequences (AAV-shCrtl) were synthesized and cloned into the GV706 vector (SM22ap-EGFP-mir155(MCS)-SV40 PolyA) using the BsmBI site (Shanghai GeneKai Technology Co., Ltd.), as detailed in Table S6. The recombinant vectors were confirmed by DNA sequencing. ApoE−/− mice were randomly divided into two groups: AAV-shTMEM41B group (n = 10); AAV-shCtrl group (n = 10). AAV-shCrtl or AAV-shTMEM41B (titer: 1E+12 vg/ml per mouse) were administered via tail vein injection. Each individual animal was considered an experimental unit, as treatments were administered to and measurements were taken from single animals.

### 2.3 Cell culturing

Human arterial smooth muscle cells (HASMC) were isolated from the arteries of healthy organ donors using established ex vivo methods, as previously described1. This procedure was approved by the Ethics Committee of the First Affiliated Hospital of Sun Yat-sen University, and donor consent was obtained. HASMC, MOVAS cells (immortalized VSMCs from mouse aortas, ATCC) and HEK293T cells (ATCC) were cultured in DMEM supplemented with 10% fetal bovine serum (Thermo Fisher), 100 U/ml penicillin, and 100 µg/ml streptomycin. Cells were maintained at 37°C in a humidified atmosphere with 95% O2 and 5% CO2.

### 2.4 Analysis of Atherosclerotic Lesions

Mice on a Western diet for a specified duration were anesthetized and euthanized. The entire aorta was excised and stained with Oil Red O (Sigma-Aldrich) for plaque analysis. The aortic root was evaluated using serial 6 µm sections. Lesion areas were measured using hematoxylin and eosin (H&E) staining. Lipid content in frozen sections was visualized with Oil Red O, while collagen deposition in lesions was assessed by Masson’s Trichrome staining. Macrophage infiltration was examined by F4/80 immunostaining.

### 2.5 Immunostaining

Frozen sections were fixed in acetone and stained following standard protocols. Cells were fixed with 4% formaldehyde, permeabilized with 0.2% Triton X-100, and blocked with 3% BSA in PBST.

Primary antibodies were applied first, followed by secondary antibodies. Imaging was performed using a Leica DMI8 or Nikon C2 confocal microscope with consistent settings.

### 2.6 Small Interfering RNA and Plasmid Transfection

For small interfering RNA (siRNA) transfection, cells at 70% confluence were transfected with gene-specific or control siRNA (50 nM), with sequences provided in Table S5, using Lipofectamine RNAi MAX Reagent (Invitrogen) according to the manufacturer’s instructions. Fresh medium was added 6 hours after transfection, and cells were incubated for 48 hours before further processing. For plasmid transfection, cells were transfected with HA-TMEM41B, FLAG-FASN, or HIS-UB overexpression plasmid, synthesized by GenePharma, at 70% confluence using Lipofectamine 3000 (Thermo Fisher) following the manufacturer’s instructions. Fresh medium was added 6 hours after transfection, and subsequent treatments were carried out at designated time points.

### 2.7 Viral infection

HSV-1 viruses used in this study were kindly provided by Prof. Ping Zhang (Zhongshan School of Medicine, Sun Yat-sen University). For in vitro experiments, cells were infected with HSV-1 at a multiplicity of infection (MOI) of 1. After culturing at 37 °C with 5% CO2 for 1h, the supernatant was discarded and then the cells were washed three times with PBS.

### 2.8 Oil-red O staining

cells exposed to oxLDL (Yiyuan Biotechnologies) for 48 hours were fixed with 4% paraformaldehyde in PBS for 15 minutes. After three PBS washes, the cells were stained with Oil Red O for 30 minutes, followed by hematoxylin staining for 2 minutes at room temperature.

### 2.9 Dil-ox-LDL uptake assay

Dil-oxLDL (Yiyuan Biotechnologies) was used to assess cholesterol uptake. Cells were treated with 10 μg/mL Dil-oxLDL for 4 hours, washed with PBS, and fixed with 4% paraformaldehyde in PBS. Cholesterol uptake was evaluated through immunofluorescence analysis.

### 2.10 Chromatin Immunoprecipitation (ChIP) Assay

OCT-1 binding sites in the TMEM41B promoter region were predicted using the JASPAR tool (http://jaspar.genereg.net/), and primers were designed to amplify the predicted binding sequence (Supplementary Table S4). ChIP assays were conducted with the Agarose ChIP Kit (Thermo Scientific Pierce). Cells were cross-linked with 1% formaldehyde for 10 minutes at room temperature, followed by quenching with glycine. DNA was immunoprecipitated from sonicated cell lysates using an OCT-1 antibody and analyzed by PCR or RT-qPCR.

### 2.11 DNA Constructs and Luciferase Reporter Assays

Cells were transfected with the appropriate plasmids in 24-well plates and harvested 48 hours later for luciferase assays. The assays were conducted using the Dual-Luciferase Reporter Assay System (Promega) following the manufacturer’s instructions. Firefly luciferase activity, normalized to Renilla luciferase, served as the internal control. Transfection experiments were performed in triplicate for each plasmid construct.

### 2.12 Immunoprecipitation (IP) and Mass spectrometry (MS)

The IP assay was conducted using the PierceTM classic magnetic IP/CO-IP kit (Thermo Fisher) according to the manufacturer’s instructions. Total proteins from HASMCs were extracted with IP lysis buffer and pre-cleared using beads. The protein complex was incubated with a specific antibody overnight at 4°C. The following day, the immunocomplex was incubated with protein A/G magnetic beads for 1 hour at room temperature. After washing to remove unbound material, the bound immune complex was eluted using a low-pH buffer. The immune complex was then subjected to mass spectrometry or Western blot analysis. For MS, peptides were extracted from SDS-PAGE gels after dehydration with 100% acetonitrile and subsequent desiccation. The peptides were analyzed by LC-MS/MS using an Easy-nLC 1200 ultra-high-performance liquid chromatography system coupled with an Orbitrap Fusion Lumos high-resolution mass spectrometer operating in data-dependent acquisition (DDA) mode.

### 2.13 Ubiquitination assay

Cells at 80% confluence were transfected with specific plasmids, followed by the indicated treatments. To prevent proteasome-mediated protein degradation, 10 μM MG132 (MCE) was added 8 hours prior to harvest. Cells were washed with ice-cold PBS, and whole-cell lysates were collected and clarified by centrifugation. The cleared lysates were used for IP with the indicated antibodies coupled to protein A/G beads (Thermo Fisher). Bound proteins were eluted using 1× SDS loading buffer, boiled, and analyzed by Western blot with the specified antibodies.

### 2.14 Statistical analysis

Data are presented as mean ± SD for normally distributed variables, or as median and IQR for non-normally distributed variables. Normality was assessed using the Shapiro-Wilk test. For normally distributed data, an unpaired two-tailed Student’s t test was used for comparisons between two groups. For multiple groups, one-way ANOVA with Bonferroni’s post hoc test was applied when variance was homogeneous. For non-normally distributed data, nonparametric tests were used: the Mann-Whitney U test for two groups and the Kruskal-Wallis test for multiple groups. Statistical analyses were performed using GraphPad Prism 8.0 software (GraphPad Software), with P < 0.05 considered statistically significant. Images without biological replicates are representative of at least three independent experiments.

## 3 Result

### 3.1 TMEM41B is Upregulated in Vascular Smooth Muscle Cells of Atherosclerotic Lesions

To elucidate the role of TMEM41B in VSMC function during AS progression, we first examined its expression in artery samples from patients with lower extremity artery diseases and healthy donors. Immunofluorescence staining revealed increased TMEM41B expression in VSMCs of AS samples (Fig. 1a-b). Western blot analysis confirmed elevated TMEM41B protein levels in AS arteries compared with normal arteries (Fig. 1c-d). To further investigate the role of TMEM41B in AS, we used apolipoprotein E–deficient (ApoE^−/−^) mice fed a Western diet (WD) and C57BL/6 mice fed a chow diet (CD) for 12 weeks. Immunofluorescence staining showed higher TMEM41B expression in VSMCs of WD-fed mice compared to CD-fed controls (Fig. 1e-f). Western blot analysis supported these findings, revealing elevated TMEM41B protein levels in the aortas of WD-fed mice (Fig. 1g-h). To mimic the atherogenic environment in vitro, primary cultured human artery smooth muscle cells (HASMCs) were stimulated with oxidized low-density lipoprotein (ox-LDL). TMEM41B protein level showed a time-dependent increase following ox-LDL treatment (Fig. 1i-j). Collectively, these findings demonstrate that TMEM41B is consistently upregulated in VSMCs under atherogenic conditions, suggesting its involvement in disease progression.

**Figure. 1.**
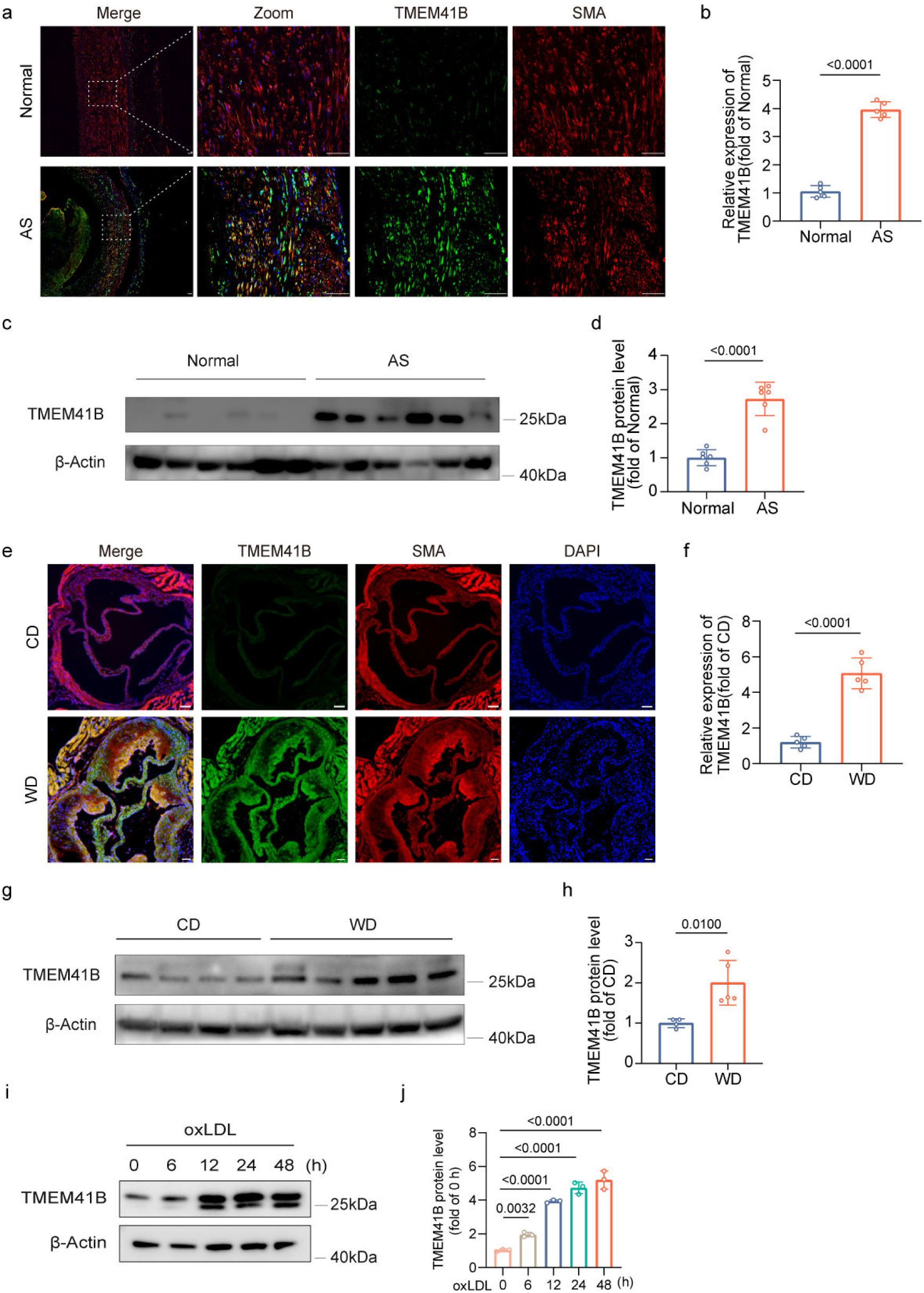
TMEM41B is Upregulated in Vascular Smooth Muscle Cells of Atherosclerotic Lesions. (a-b) Representative immunofluorescence images showing co-localization of TMEM41B with α-SMA, a vascular smooth muscle cell (VSMC)-specific marker, in normal and atherosclerotic (AS) human arteries (n = 5 per group). Nuclei were counterstained with DAPI. Scale bar: 50 μm. (c-d) Western blot analysis and quantification showing increased TMEM41B protein levels in AS human arteries compared to normal controls (n = 6 per group). (e-f) Representative immunofluorescent images of TMEM41B and α-SMA expression in aortas from C57BL/6 mice fed a chow diet (CD) or Apoe^−/−^ mice fed a Western diet (WD) for 12 weeks (n = 5 per group). Scale bar: 20 μm. (g-h) Western blot analysis and quantification of TMEM41B protein expression in aortas from CD-fed C57BL/6 mice (n = 4) and WD-fed Apoe^−/−^ mice (n = 5). (i-j) Western blot analysis and quantification of TMEM41B protein levels in human artery smooth muscle cells (HASMCs) treated with oxidized low-density lipoprotein (oxLDL, 50 μg/mL) for the indicated time points (n = 3). (a-h) Data were analyzed using an unpaired, two-tailed Student’s t-test. (i-j) Data were analyzed using one-way ANOVA.

### 3.2 VSMC-Specific TMEM41B Silencing Reduces Atherosclerosis in ApoE**^−/−^** Mice

To investigate the potential role of TMEM41B in atherosclerosis, we employed AAV9 vectors with the VSMC-specific promoter SM22α to knock down TMEM41B in the VSMCs of ApoE^−/−^ mice (Fig. 2a). This approach achieved efficient and specific knockdown of TMEM41B in VSMCs, with no detectable off-target effects in other tissues (Fig. S1). After 14 weeks on a WD, en face Oil Red O staining demonstrated a clear reduction in atherosclerotic plaque size in AAV-shTMEM41B mice compared to controls (Fig. 2b). We then conducted a detailed analysis of the aortic root components (Fig. 2c-d). Histological analysis of aortic root sections revealed that VSMCs specific TMEM41B knockdown reduced atherosclerotic lesion area and lipid content, confirmed by HE and Oil Red O staining. Furthermore, Masson’s trichrome staining indicated an increase in collagen content after silencing TMEM41b expression. Immunofluorescence analysis also showed reduced infiltration of F4/80^+^ macrophages in plaques, suggesting that TMEM41B knockdown limits inflammatory activity while promoting features of plaque stabilization. Importantly, no significant differences were observed in body weight or serum lipid levels, including triglycerides (TG), total cholesterol (TC), LDL-C, and HDL-C, between AAV-shTMEM41B and control groups (Fig. S2, Fig. 2e). This indicates that the observed reduction in atherosclerosis is independent of systemic metabolic changes. These findings identify TMEM41B as a critical regulator of VSMC-driven atherosclerosis, demonstrating that its targeted knockdown effectively attenuates AS progression.

**Figure. 2.**
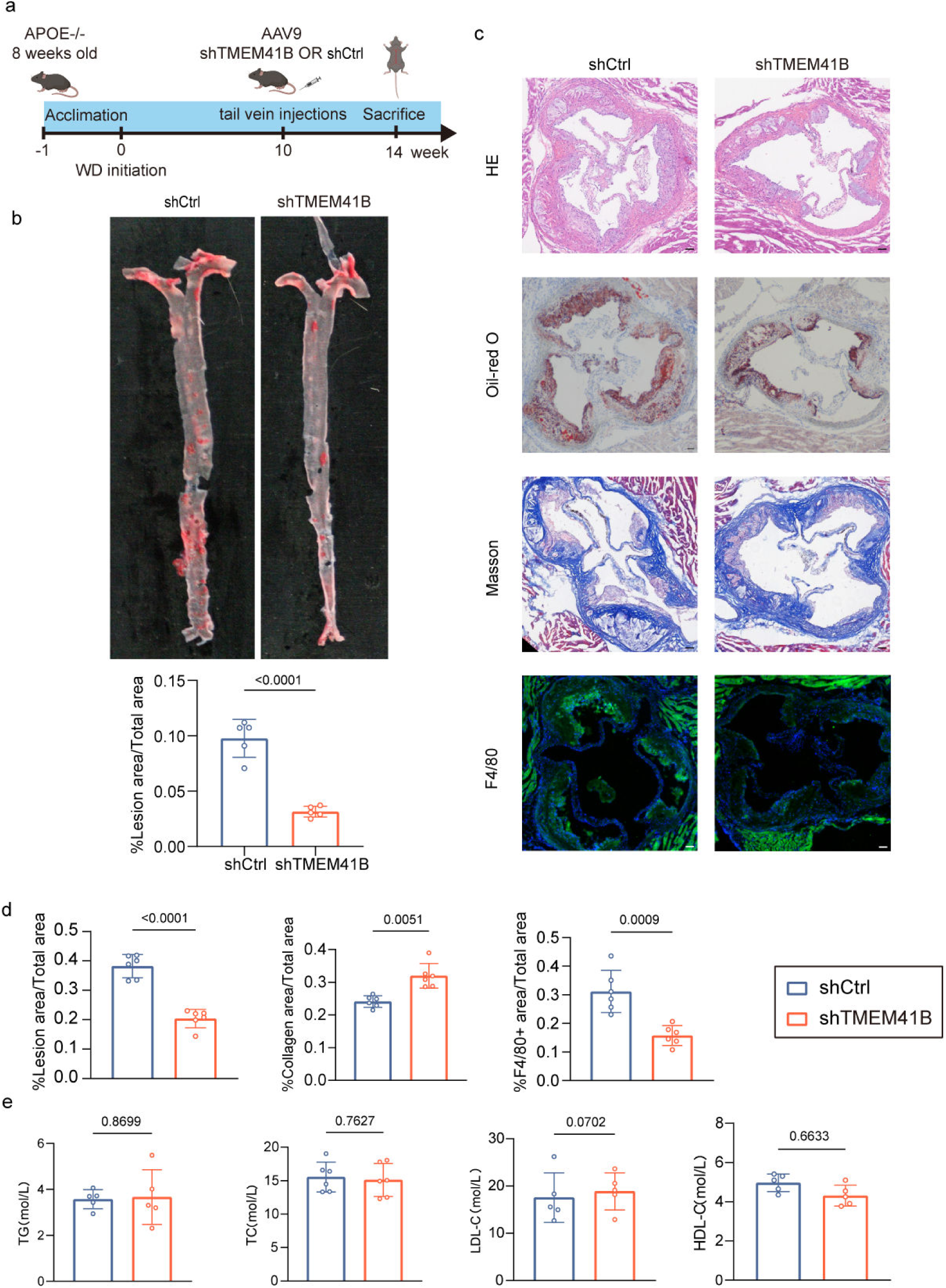
VSMC-Specific TMEM41B Silencing Reduces Atherosclerosis in Apoe^−/−^ Mice. (a) Schematic representation of the experimental protocol for atherosclerosis assessment in ApoE^−/−^ mice with VSMC-specific TMEM41B knockdown. (b) En face Oil Red O staining of aortas and quantification from AAV-shTMEM41B mice in comparison to controls (n = 5). (c) Representative images of aortic root sections stained for lesion area (H&E), lipid content (Oil Red O), collagen deposition (Masson’s trichrome), and macrophage infiltration (F4/80). Scale bar: 20 μm (n = 5). (d) Quantification of lesion area, collagen content, and F4/80^+^ areas in plaques (n = 5). (e) Serum lipid levels, including triglycerides (TG), total cholesterol (TC), LDL-C, and HDL-C, in control and AAV-shTMEM41B groups (n = 5 per group). (a-e) Data were analyzed using an unpaired, two-tailed Student’s t-test.

### 3.3 TMEM41B Modulates Lipid Profiles in VSMCs

To investigate how VSMC-specific TMEM41B deficiency attenuates atherosclerosis, we conducted RNA-Seq on TMEM41B-knockdown and TMEM41B-overexpressing mouse aortic vascular smooth muscle cells (MOVAS). Kyoto Encyclopedia of Genes and Genomes (KEGG) and Gene Ontology (GO) enrichment analyses revealed that differentially expressed genes were significantly enriched in pathways related to lipid metabolism and inflammation (Fig. 3a). Gene Set Enrichment Analysis (GSEA) further highlighted fatty acid metabolism and fatty acid biosynthetic process as among the most significantly upregulated pathways associated with TMEM41B expression (Fig. 3b, Fig. S3). We hypothesize that TMEM41B deficiency alleviates atherosclerosis by regulating VSMCs lipid metabolism, thereby reducing the formation of VSMC-derived foam cells.

**Figure. 3.**
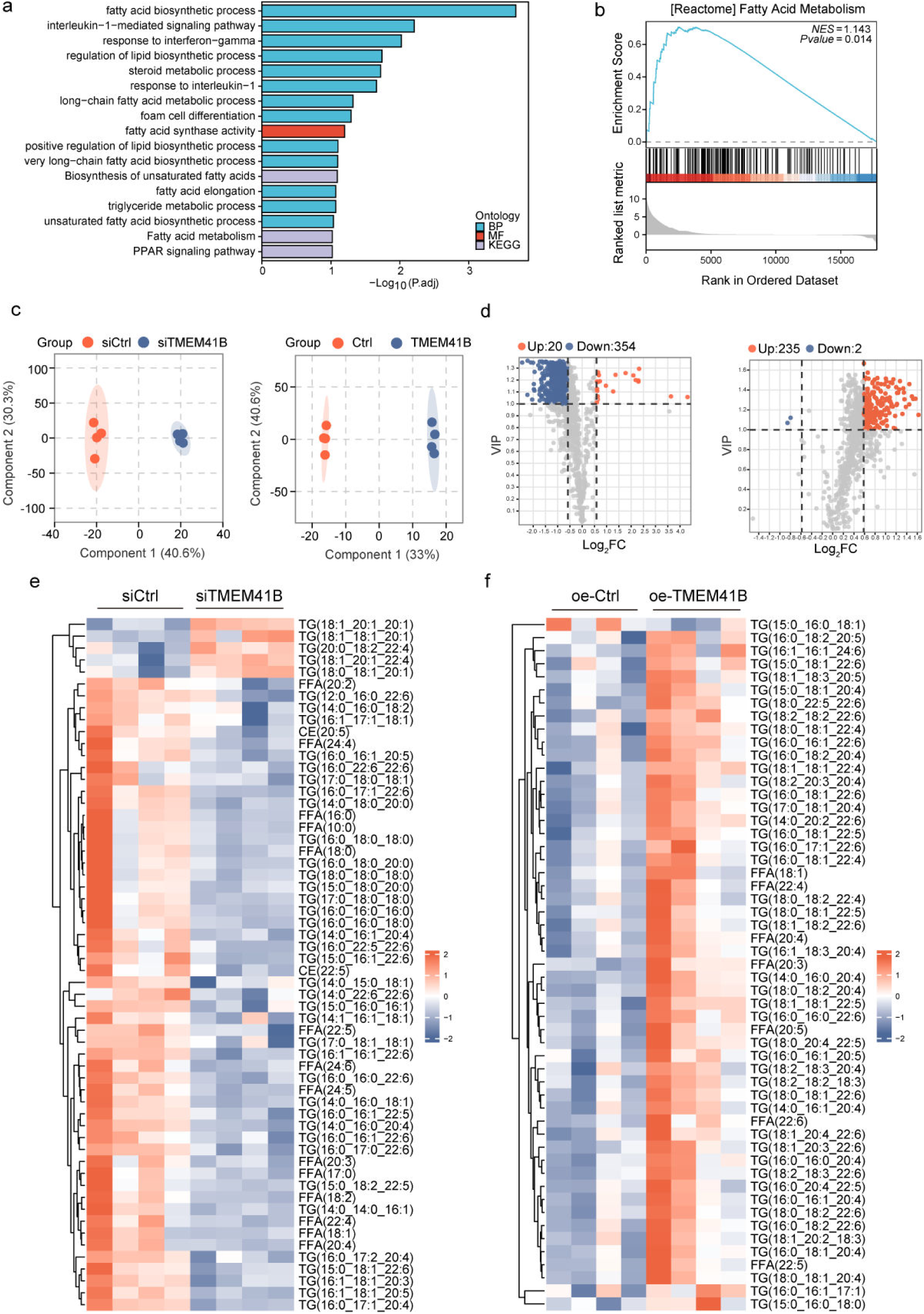
TMEM41B Modulates Lipid Profiles in VSMCs. (a) Gene Ontology (GO) and Kyoto Encyclopedia of Genes and Genomes (KEGG) pathway enrichment analyses of differentially expressed genes (DEGs) between TMEM41B-overexpressing (oe-TMEM41B) and control (oe-Ctrl) MOVAS. (b) Gene set enrichment analysis (GSEA) highlighting the enrichment of fatty acid metabolism pathways in oe-TMEM41B compared with oe-Ctrl group. (c-e) HASMCs were transfected with siCtrl/siTMEM41B or oe-Ctrl/oe-TMEM41B and subsequently analyzed using lipidomic assays (n=4 per group). (c) Orthogonal partial least squares discriminant analysis (OPLS-DA) plots demonstrating lipidomic variations across different groups. (d) Volcano plots visualizing differentially abundant metabolites across groups, with significant thresholds defined as VIP ≥ 1 and |log_2_FC| ≥ 0.58. (e) Heatmap illustrating changes in triglycerides (TGs), cholesteryl esters (CEs), and free fatty acids (FFAs) levels across experimental conditions.

To validate this hypothesis, we performed a comprehensive lipidomics analysis. Orthogonal Partial Least Squares Discriminant Analysis (OPLS-DA) models demonstrated that TMEM41B knockdown and overexpression induced significant shifts in intracellular lipid profiles (Fig. 3c). Specifically, TMEM41B knockdown reduced levels of 422 lipid species, while increasing 21 lipid species. In contrast, TMEM41B overexpression upregulated 232 lipids while downregulating only 2, indicating a strong impact on cellular lipid composition (Fig. 3d).

Previous studies have identified triglycerides, cholesterol esters, and non-esterified fatty acids as key lipid components in VSMC-derived foam cells, all of which are known to promote atherosclerosis[28–30]. Heatmap analysis demonstrated a significant reduction in triglycerides, cholesterol, and non-esterified fatty acid levels in TMEM41B knockdown HASMCs compared to controls, while these lipid levels were notably elevated in TMEM41B-overexpressing HASMCs (Fig. 3e-f). Our findings extend prior studies by demonstrating that TMEM41B plays a pivotal role in modulating lipid metabolism in VSMCs, which contributes to foam cell formation.

### 3.4 TMEM41B Promotes VSMC-derived Foam Cell Formation

Foam cell formation is a critical process in the pathogenesis of atherosclerosis. ox-LDL exposure enhances foam cell formation and modulates inflammatory responses in VSMCs[31, 32]. To determine the role of TMEM41B in VSMC-derived foam cell formation, HASMCs were treated with ox-LDL for 48 hours following siRNA-mediated knockdown or plasmid-mediated overexpression of TMEM41B to simulate foam cell formation in vitro. Oil Red O staining revealed that TMEM41B overexpression markedly increased lipid droplet formation in HASMCs, while this process was significantly attenuated by TMEM41b knockdown (Fig. 4a-b). A cholesterol uptake assay using Dil-oxLDL incubation demonstrated that TMEM41B did not alter cholesterol uptake in HASMCs (Fig. 4c-d). Consistently, TMEM41B overexpression significantly increased intracellular triglyceride (TG) and total cholesterol (TC) levels, whereas knockdown produced the opposite effect (Fig. 4e-f). We further analyzed the expression of key proteins involved in lipid uptake, synthesis, and cholesterol efflux pathways. Western blot showed that overexpressing TMEM41B in HASMCs obviously increased Fatty Acid Synthase (FASN, key enzyme in fatty acid synthesis) protein level, while silencing TMEM41b significantly reduced FASN expression, with no changes observed in scavenger receptors (CD36, LOX-1, SRA) or the cholesterol efflux transporter (ABCA1) (Fig. 4g-j). As lipid metabolism and inflammation are tightly interconnected in atherosclerosis, we next examined the inflammatory phenotype of HASMCs. TMEM41B overexpression significantly upregulated pro-inflammatory cytokines IL1β, IL6 and TNFα, whereas TMEM41b knockdown reduced their level (Fig. S4). This indicates that TMEM41B not only modulates lipid accumulation but also drives inflammatory responses in VSMC-derived foam cells. In summary, these findings suggest that TMEM41B enhances VSMC-derived foam cell formation during the process of atherosclerosis.

**Figure. 4.**
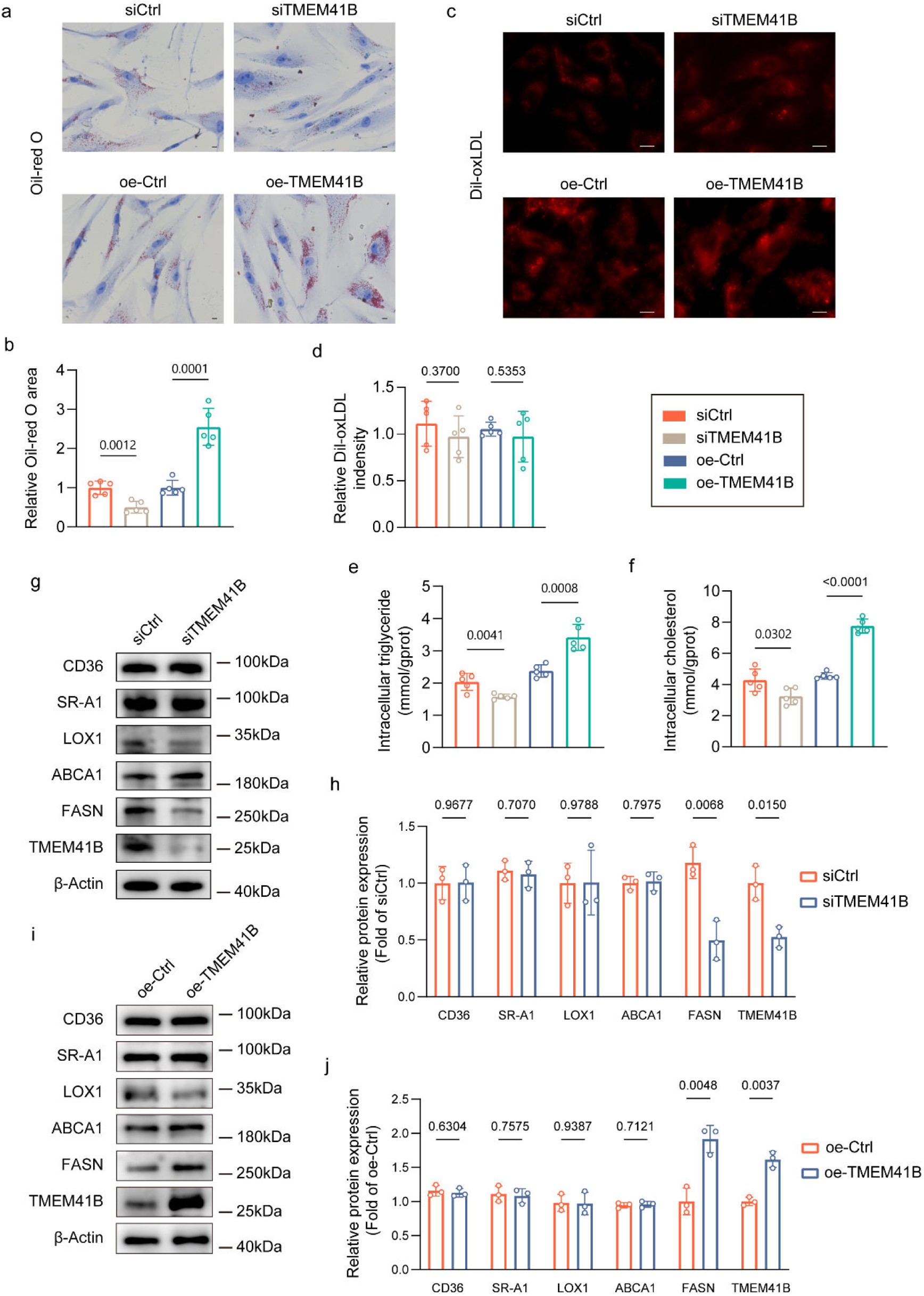
TMEM41B Promotes VSMC-derived Foam Cell Formation. (a-b) Oil Red O staining of HASMCs transfected with siCtrl or siTMEM41B (upper panel) and oe-Ctrl or oe-TMEM41B (lower panel) and incubated with oxidized LDL (ox-LDL, 50 μg/mL) for 48 hours. Scale bar: 20 μm. (c-d) Immunofluorescence analysis of HASMCs transfected with siCtrl/siTMEM41B (upper panel) or oe-Ctrl/oe-TMEM41B (lower panel) and treated with Dil-oxLDL (30 μg/mL) for 4 hours. Quantification of Dil-ox-LDL fluorescence intensity. Scale bar 20um. (e-f) Intracellular triglyceride (TG) and total cholesterol (TC) levels in HASMCs following TMEM41B knockdown or overexpression. (n = 5). (g-h) Western blot analysis and quantification of indicated proteins in HASMCs transfected with siCtrl or siTMEM41B. (i-j) Western blot analysis and quantification of indicated proteins in HASMCs transfected with oe-Ctrl or oe-TMEM41B. (a-j) Data were analyzed using an unpaired, two-tailed Student’s t-test.

### 3.5 TMEM41B Directly Binds FASN and Inhibits its Ubiquitination and Proteasomal Degradation

To elucidate the mechanism of TMEM41B in VSMCs derived foam cell formation, we conducted immunoprecipitation coupled with mass spectrometry (IP/MS) to identify TMEM41B-interacting proteins. Among the top 10 interactors identified, FASN was prominently featured (Fig. S5). This finding aligns with our prior observations linking TMEM41B to lipid metabolism. To validate the TMEM41B-FASN interaction, HASMCs were transfected with HA-TMEM41B or FLAG-FASN constructs, followed by reciprocal IP assays using anti-HA antibodies or anti-FLAG antibodies. Immunoblotting confirmed that TMEM41B interacted with FASN (Fig. 5a-b). Immunofluorescence staining further demonstrated co-localization of TMEM41B and FASN in HASMCs and MOVAS (Fig. 5c and Fig. S6). We additionally mapped the interaction sites between TMEM41B and FASN through computational protein-protein interaction (PPI) analysis using 3D models of both proteins. The AutoDock algorithm was employed to predict the highest-confidence interaction pose (Figure. 5d).

**Figure. 5.**
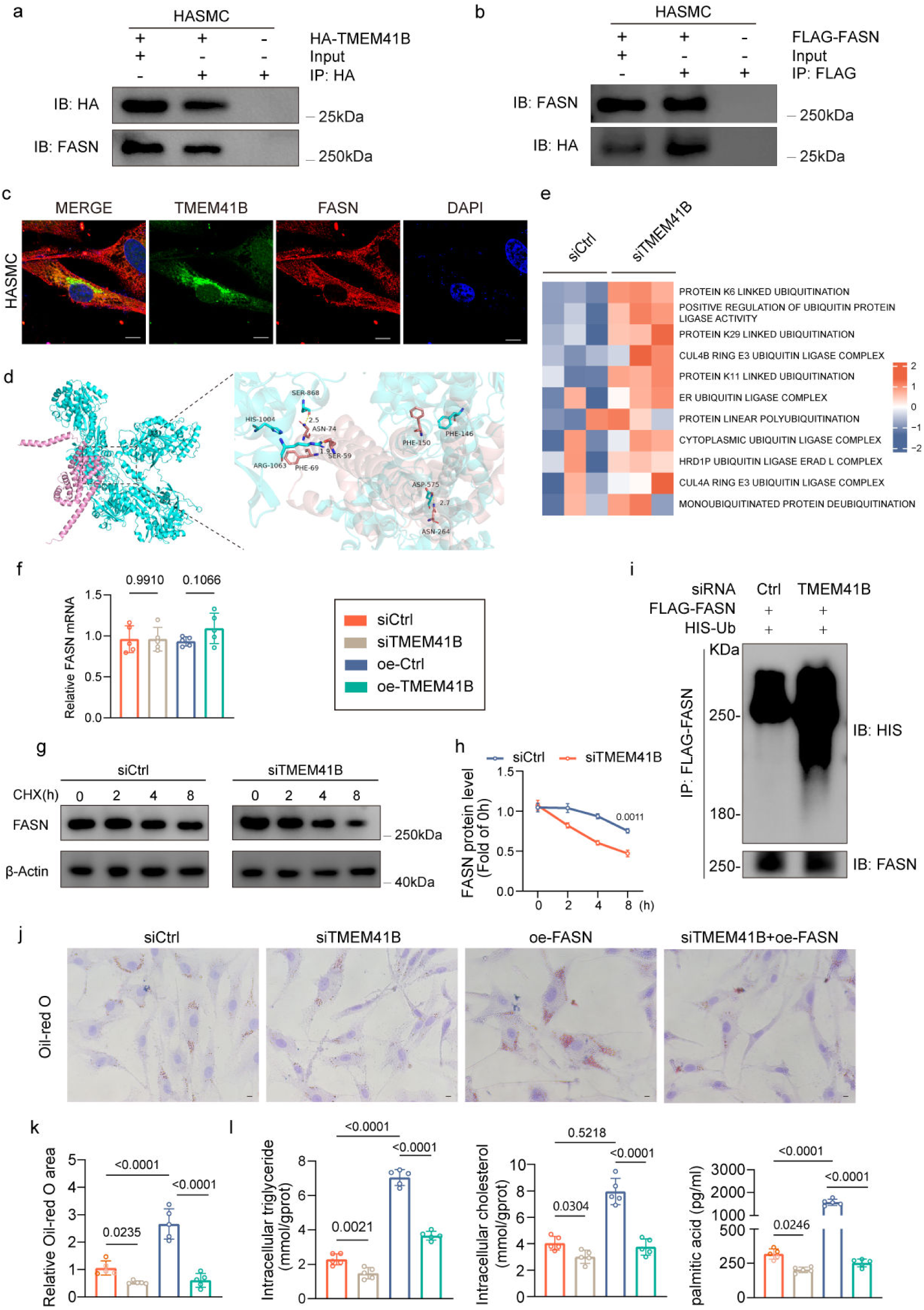
TMEM41B Directly Binds FASN and Inhibits its Ubiquitination and Proteasomal Degradation. (a-b) HASMCs were transfected HA-TMEM41B, FLAG-FASN or empty vector, after 48 hours of transfection, Immunoprecipitation (IP) and immunoblotting (IB) were performed with the indicated antibodies. (c) HASMCs were subjected to immunofluorescent staining with an anti-TMEM41B (green) or anti-FASN (red) antibody. Nuclei were counterstained with DAPI. Scale bar: 10 μm. (d) Molecular docking analysis illustrating the interaction between TMEM41B and FASN. (e) GSVA analysis of ubiquitin signaling pathways between siCtrl and siTMEM41B groups. (f) RT-qPCR analysis of FASN gene expression in HASMCs following TMEM41B knockdown or overexpression. (g-h) Western blot analysis and quantification showing FASN protein stability in HASMCs treated with cycloheximide (50 μg/mL) at the indicated time points. (i) Western blot analysis of indicated proteins in HASMCs cotransfected with flag-FASN and His-Ub in the presence of siCtrl or siTMEM41B, with proteasome inhibitor MG132 (10 μM) treatment for 8 hours before IP of whole cell lysates with Flag magnetic beads (n = 3). (j-k) Oil Red O staining and quantification of lipid accumulation in HASMCs transfected with siCtrl, siTMEM41B, oe-FASN or cotransfected with siTMEM41B and oe-FASN, treated with oxLDL (50 μg/mL) for 48 hours (n=5). Scale bar: 20 μm. (l) Quantitative intracellular triglyceride (TG), total cholesterol (TC) and palmitic acid levels in HASMCs transfected with siCtrl, siTMEM41B, oe-FASN or cotransfected with siTMEM41B and oe-FASN (n = 5). (f) Data were analyzed using an unpaired, two-tailed Student’s t-test, (g-h) two-way ANOVA, (j-l) one-way ANOVA.

Interestingly, Gene set variation analysis (GSVA) analysis revealed that ubiquitin-related pathway activity was upregulated in the TMEM41B knockdown group, whereas it was significantly suppressed in the overexpression group (Fig. 5e, Fig. S7). Based on these findings, we hypothesize that TMEM41B regulates foam cell formation by modulating FASN ubiquitination levels. As previously shown, TMEM41B knockdown reduced FASN protein expression, whereas its overexpression increased FASN levels. Notably, FASN mRNA levels remained unchanged, indicating that TMEM41B regulates FASN through post-translational mechanisms (Fig. 5f). In support of this notion, cycloheximide (CHX)-chase assays showed that FASN stability decreased in TMEM41B knockdown cells (Fig. 5g-h). To assess whether TMEM41B modulates FASN ubiquitination, we overexpressed FLAG-FASN and His-ubiquitin in HASMCs cells transfected with either control or TMEM41B siRNA. Immunoblotting revealed elevated FASN ubiquitination in TMEM41B knockdown cells relative to controls (Figure. 5i). These findings suggest that TMEM41B directly interacts with FASN and regulates its levels via the ubiquitin-proteasome pathway.

To assess the interplay between TMEM41B and FASN in foam cell formation, we concurrently knocked down TMEM41B and overexpressed FASN in HASMCs. Oil Red O staining (Fig. 5j-k) and quantification of intracellular TG, TC and palmitate—a key fatty acid product of FASN activity-demonstrated that FASN overexpression significantly promoted foam cell formation, resulting in increased neutral lipid accumulation (Fig. 5l). However, TMEM41B deficiency effectively mitigated these effects These findings highlight the critical role of the TMEM41B/FASN axis in modulating VSMC-derived foam cell formation, providing novel insights into the regulation of lipid metabolism and its implications for atherosclerosis progression.

### 3.6 HSV Infection Enhances TMEM41B Expression via OCT-1-Mediated Transcriptional Activation

Epidemiological studies have linked HSV infection to an elevated risk of atherosclerosis[33, 34], and the presence of HSV in atherosclerotic lesions has been established[33]. TMEM41B is a host factor required for diverse virus replication[21, 26]. Interestingly, a significant increase in both TMEM41B mRNA and protein levels was observed in HASMCs following HSV infection (Fig. 6a-b). Concurrently, Oil-red O staining revealed enhanced lipid droplets accumulation in HSV-infected HASMCs, while this effect was effectively reversed by TMEM41B knockdown (Fig. 6c). Given that OCT-1 is a key host transcription factor activated during HSV infection[35], we hypothesized that HSV promotes TMEM41B transcription via OCT-1. Promoter analysis with the JASPAR database identified multiple putative OCT-1 binding sites within the TMEM41B promoter region.

**Figure. 6.**
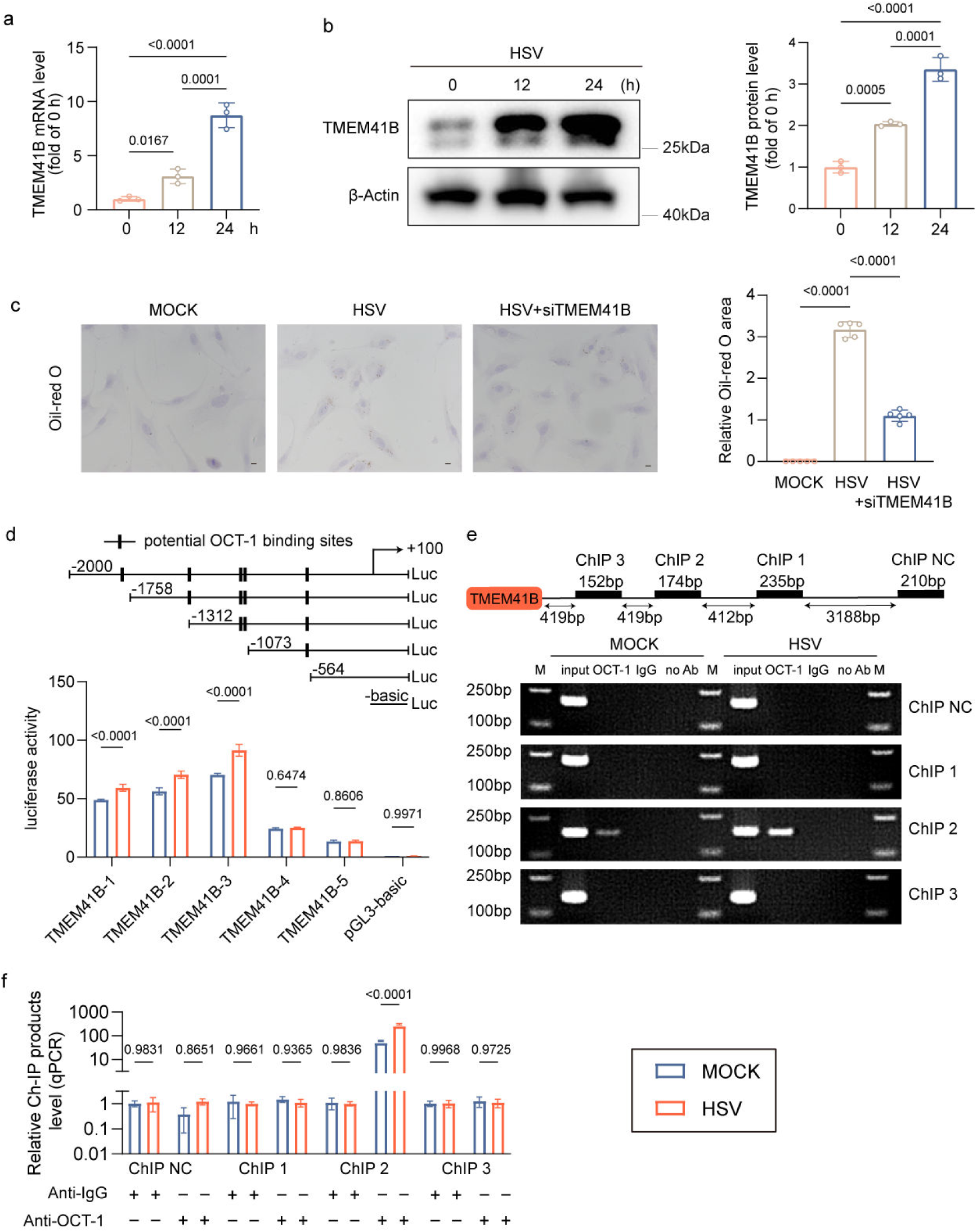
HSV Infection Enhances TMEM41B Expression via OCT-1-Mediated Transcriptional Activation. (a-b) RT-qPCR and western blot analysis showing increased TMEM41B mRNA and protein levels in HASMCs infected with HSV (MOI = 1) for the indicated times (n = 3). (c) Oil Red O staining and quantification of lipid accumulation in HASMCs transfected with siCtrl or siTMEM41B following HSV infection for 24 hours (MOI = 1) (n = 5). Scale bar: 20 μm. (d) Schematic representation of truncated constructs of the TMEM41B promoter reporter. Luciferase reporter assays showing TMEM41B promoter activity in mock– and HSV-infected 293T cells. (e) Chromatin immunoprecipitation (ChIP) assay validating OCT-1 binding to the TMEM41B promoter, with enhanced binding following HSV infection in HASMCs. M: Marker. (f) RT-qPCR of the Ch-IP products validated the binding capacity of OCT-1 to the TMEM41B promoter. (a-c) Data were analyzed using the one-way ANOVA, (d-f) two-way ANOVA.

To further investigate the mechanism underlying HSV-induced transcriptional upregulation of TMEM41B, we constructed truncated variants of the full-length TMEM41B promoter and transfected them into 293T cells, followed by HSV treatment. This analysis revealed that HSV-induced TMEM41B regulation was mediated by potential OCT-1 binding sites located between −2000 and −1073 bp (Fig. 6d). To further validate the involvement of OCT-1, chromatin immunoprecipitation (ChIP) assays confirmed direct OCT-1 binding to a specific segment of the TMEM41B promoter between −1315 and −1141 bp, which was further enhanced following HSV infection (Fig. 6 e-f). In summary, our results reveal that HSV infection activates OCT-1, which directly enhances TMEM41B transcription. This leads to increased lipid accumulation in HASMCs and highlighting a novel link between viral infection and metabolic reprogramming in atherosclerosis.

## 4 Discussion

Disturbed lipid metabolism and the inflammatory response within plaques are key drivers of AS progression, with foam cells playing a central role in this process. However, the exact mechanisms underlying their involvement remain unclear. Here, we identify a novel regulatory mechanism in VSMC-derived foam cell formation. Specifically, our findings reveal that TMEM41B was markedly upregulated in both human and murine atherosclerotic lesions. In cultured VSMCs, TMEM41B promotes foam cell formation by enhancing lipid synthesis and the secretion of inflammatory cytokines. Using VSMC-specific TMEM41B silencing in Aope^−/−^ mice, we confirmed that TMEM41B inhibition suppresses atherosclerosis progression. Mechanistically, TMEM41B directly interacts with FASN, inhibiting its ubiquitination and degradation, thereby promoting VSMC lipid accumulation. Additionally, we uncover a novel link between HSV infection and AS progression through OCT-1-mediated upregulation of TMEM41B, revealing a critical interplay between viral infection and metabolic reprogramming in vascular pathologies. These findings highlight the pathogenic role of lipid-laden VSMCs in atherosclerotic plaques and suggest that TMEM41B may serve as a new therapeutic target for atherosclerosis.

Traditionally, macrophages in atherosclerotic plaques are believed to arise from the recruitment of monocytes from the bloodstream and the proliferation of resident tissue macrophages[35]. These macrophages uptake oxidized low-density lipoproteins (oxLDL), transforming into foam cells, a key pathological feature of atherosclerosis[9]. Foam cells secrete both pro-inflammatory and anti-inflammatory cytokines, recruit additional immune cells, and clear apoptotic cells[36]. Recent advances in single-cell sequencing and lineage tracing have demonstrated that VSMCs and their derivatives are the primary cellular contributors to the development of atherosclerotic lesions at various stages[37–39]. Notably, more than 50% of foam cells in these lesions originate from VSMCs, rather than from circulating monocytes[11, 12]. In healthy vessels, vascular smooth muscle cells (VSMCs) maintain a contractile phenotype. However, during atherosclerosis, VSMCs become activated and transition to a proliferative or synthetic phenotype, marked by increased cell division and protein synthesis[40]. In this study, we identify TMEM41B as a critical regulator of VSMC-derived foam cell formation. By silencing TMEM41B in VSMCs both in vitro and in vivo, we observed a significant reduction in foam cell formation and a decrease in plaque development. These results suggest that targeting VSMC-derived foam cells could be an effective strategy to suppress plaque formation. This is the first study to link TMEM41B to foam cell formation in VSMCs, challenging the traditional macrophage-centric view of atherosclerosis.

While statins have significantly reduced cardiovascular risk, genetic variations and metabolic differences contribute to residual risk in some patients[41, 42]. Studies show that approximately 50% of patients do not achieve the expected benefits from statin therapy[43]. Recent advances in lipidomics have highlighted substantial differences in lipid compositions within atherosclerotic plaques[44, 45] The key components of plaques, foam cells, play a pivotal role in lipid metabolic remodeling. Understanding the metabolic characteristics of foam cells could lead to more effective strategies for combating atherosclerosis. Previous studies have reported lipidomic features of macrophage-derived foam cells, including elevated levels of oxidized sterols, sphingolipids, and ceramides in human atherosclerotic plaques[46]. Interestingly, these lipid species were significantly reduced upon engagement with cholesterol receptors, particularly apoA-I. Early studies have shown that VSMCs accumulate triglycerides under high-lipid conditions, highlighting their adaptability to lipid fluctuations and reflects their metabolic adaptability to pathological conditions[47, 48]. In line with these findings, our study is the first to examine the lipidomic profile of VSMC-derived foam cells. Through lipidomic analysis, we demonstrate that TMEM41B regulates lipid metabolism reprogramming in VSMCs. Silencing TMEM41B resulted in a substantial reduction in triglycerides, cholesterol, long-chain fatty acids, and ceramides—lipids known to promote atherosclerosis. These results suggest that TMEM41B-driven lipid reprogramming in VSMCs is a key mechanism in the progression of VSMC-derived foam cells.

Despite progress in understanding lipid metabolism in atherosclerosis, the mechanisms underlying lipid reprogramming in VSMCs remain poorly defined. Recent studies on VSMC-derived foam cells have identified mechanisms such as defective autophagy and impaired reverse cholesterol transport, including reduced ABCA1 expression, as contributing factors to lipid accumulation in VSMCs[11, 16, 17]. Interestingly, one study found that VSMC-derived foam cells exhibit lower phagocytic and clearance capacities compared to macrophage-derived foam cells and show a weaker immune response[49]. This suggests that VSMCs may have distinct lipid metabolic profiles compared to macrophages. Further research has revealed that, in the early stages of atherosclerotic lesions, VSMCs in diffuse intimal thickening (DIT) possess more synthetic organelles—such as the rough endoplasmic reticulum, ribosomes, and mitochondria—compared to medial VSMCs[50]. De-differentiated VSMCs also increase extracellular matrix (ECM) synthesis and secrete more pro-inflammatory cytokines[51]. These findings suggest that VSMCs may upregulate their lipid synthesis capacity under atherosclerotic conditions. Our findings represent a significant shift in understanding lipid accumulation in VSMC-derived foam cell. We identify a novel mechanism by which VSMCs promote foam cell formation by increasing lipid synthesis, rather than by altering lipid uptake or efflux. Through an IP-MS screening, we identified FASN as a direct interacting partner of TMEM41B. Further co-immunoprecipitation (Co-IP) and ubiquitination experiments demonstrated that TMEM41B acts as a stabilizer of FASN, a key enzyme in fatty acid synthesis[52]. Notably, TMEM41B upregulates FASN expression and enhances fatty acid synthesis in VSMCs, further corroborating our hypothesis. These findings offer a novel insight into the lipid metabolic reprogramming of VSMCs and provide potential new therapeutic targets for atherosclerosis.

Post-translational modifications such as ubiquitination regulate FASN activity and are implicated in lipid dysregulation and metabolic diseases. For instance, TRIM56 promotes fatty liver by modulating FASN ubiquitination[53], while USP14 stabilizes FASN via deubiquitination and contributes to hepatic steatosis [54]. Consistent with this, our present data revealed that TMEM41B overexpression suppresses ubiquitin-related pathways, while its knockdown leads to their upregulation. Although previous studies linked TMEM41B to the regulation of autophagy and phospholipid scramblase activity, our findings uncover its unique role in stabilizing FASN to promote foam cell formation. However, the specific E3 ligases or deubiquitinases involved in TMEM41B-mediated regulation remain unidentified. Future studies are needed to elucidate the precise mechanisms by which TMEM41B mediates FASN ubiquitination.

FASN-mediated fatty acid synthesis is central to linking lipid metabolism with inflammatory signaling and disease progression. For instance, FASN has been shown to promote atherosclerosis by regulating fatty acid synthesis pathways[48]. Recent studies further demonstrate that FASN contributes to foam cell formation through cholesterol-induced KLF4 upregulation and transformation of VSMCs[55]. Additionally, FASN also drives cholesterol synthesis by promoting acetoacetyl-CoA production[56]. This metabolite plays a critical role in regulating crosstalk between cholesterol and inflammatory signaling. Such metabolic reprogramming likely amplifies inflammatory responses by generating pro-inflammatory lipid mediators. Building on these findings, our data demonstrate that TMEM41B overexpression in VSMCs led to increased secretion of pro-inflammatory cytokines, including IL-1β, IL-6, and TNF-α. While the exact mechanism remains unclear, lipid intermediates produced by the TMEM41B/FASN axis may act as signaling molecules to activate inflammatory pathways. Further investigation is warranted to elucidate the precise mechanisms linking TMEM41B to inflammatory responses and lipid metabolism.

TMEM41B is also a critical player in viral replication, particularly for RNA viruses like SARS-CoV-2 and flaviviruses[21, 26]. These viruses create replication compartments within the endoplasmic reticulum membrane, and TMEM41B has been identified as an essential host factor for this process. Epidemiological studies have established a link between HSV infection and increased AS risk[33, 34], highlighting the importance of uncovering the underlying mechanisms. Expanding upon these observations, our study provides mechanistic insights into how HSV infection promotes foam cell formation through metabolic reprogramming. Specifically, we found that HSV infection upregulates TMEM41B expression through OCT-1 activation. Preliminary ChIP-PCR experiments support this mechanism, indicating that OCT-1 directly binds to the TMEM41B promoter region. This pathway likely accelerates foam cell formation by enhancing lipid synthesis, highlighting the role of TMEM41B in linking viral infection with AS progression. Prior research has shown that OCT-1 plays a dual role as a transcriptional regulator in both AS[57, 58] and HSV infection[59, 60]. During the early stages of infection, OCT-1 supports viral gene expression while broadly altering the host cell transcriptome, potentially reprogramming lipid metabolism to favor foam cell formation. Our study further elucidates the regulatory relationship between OCT-1 and TMEM41B, providing new insights into the interplay between viral infection and metabolic dysregulation in vascular disease. While the precise molecular mechanisms remain to be clarified, our findings suggest that targeting the OCT-1/TMEM41B axis could offer therapeutic potential, particularly for preventing AS progression or inhibiting plaque formation in HSV-infected patients. Future studies should explore whether OCT-1-mediated TMEM41B upregulation is specific to HSV infection or represents a broader host response to viral infections. Additionally, investigating the involvement of chromatin remodeling factors or co-activators in this regulatory pathway could further illuminate the molecular underpinnings of this novel mechanism.

In summary, our study establishes TMEM41B as a central regulator of foam cell formation. By stabilizing FASN, TMEM41B enhances fatty acid synthesis and drives pro-inflammatory lipid mediator production, shedding light on the metabolic underpinnings of atherosclerotic plaque development. These findings redefine the paradigm of foam cell formation, shifting focus from macrophage-dominated mechanisms to VSMC-driven processes. Targeting TMEM41B could allow for precise modulation of foam cell formation and inflammatory responses, particularly in patients with AS linked to viral infections such as HSV.

## Acknowledgements

We thank Prof. Ping Zhang (Zhongshan School of Medicine, Sun Yat-sen University) for providing HSV-1 viruses and technical assistance.

