## Supplementary material for "TMEM41B Contributes to Atherosclerosis by Promoting Lipid Synthesis in Vascular Smooth Muscle Cells via Fatty Acid Synthase Stabilization": Supplementary Information.docx

1. Supplementary methods

1.1 Mass spectrometry-based lipidomics

Samples were removed from the −80°C freezer and thawed on ice. An extraction solvent (MTBE:MeOH = 3:1, v/v) containing an internal standard mixture (1 mL) was added to each sample. The mixture was vortexed for 15 minutes, followed by the addition of 200 μL of water. After vortexing for 1 minute, the samples were centrifuged at 12,000 rpm for 10 minutes. A 200 μL aliquot of the upper organic layer was collected and evaporated using a vacuum concentrator. The dried extracts were reconstituted in 200 μL of solution (ACN:IPA = 1:1, v/v) for LC-MS/MS analysis.

The extracts were analyzed using an LC-ESI-MS/MS system (UPLC, ExionLC AD; MS, QTRAP® System, SCIEX). UPLC analysis utilized a Thermo Accucore™ C30 column (2.6 μm, 2.1 mm × 100 mm). The solvent system consisted of A: acetonitrile/water (60/40, v/v, 0.1% formic acid, 10 mmol/L ammonium formate) and B: acetonitrile/isopropanol (10/90, v/v, 0.1% formic acid, 10 mmol/L ammonium formate). The gradient program was as follows: A/B (80:20, v/v) at 0 min, 70:30 at 2.0 min, 40:60 at 4 min, 15:85 at 9 min, 10:90 at 14 min, 5:95 at 15.5 min, maintained at 5:95 until 17.3 min, then reset to 80:20 at 17.3 min and held until 20 min. The flow rate was 0.35 mL/min, column temperature was 45°C, and the injection volume was 2 μL. The effluent was connected to an ESI-triple quadrupole-linear ion trap (QTRAP)-MS system operating in positive and negative ion modes, controlled by Analyst 1.6.3 software (SCIEX). Key source parameters included: turbo spray ion source, source temperature of 500°C, ion spray voltage at 5500 V (positive mode) and −4500 V (negative mode), and gas pressures (GS1, GS2, CUR) set to 45, 55, and 35 psi, respectively. The collision gas (CAD) was set to medium. Instrument tuning and calibration were performed with 10 and 100 μmol/L polypropylene glycol solutions in QQQ and LIT modes. MRM transitions were optimized for DP and CE, with a specific set of transitions monitored based on the elution period of the metabolites.

1.2 RNA-sequencing and gene expression analysis

Total RNA was extracted from MOVAS under specific conditions using Trizol reagent (Thermo Fisher). RNA sequencing (RNA-seq) was performed by BGI-Shenzhen Company following their standard protocols. Sequencing data underwent quality control using SOAPnuke, which included: (1) removal of reads with sequencing adapters, (2) exclusion of reads with a low-quality base ratio (base quality ≤ 15) exceeding 20%, and (3) elimination of reads containing an unknown base ('N') ratio greater than 5%. High-quality clean reads were generated and stored in FASTQ format for further analysis. Clean reads were mapped to the reference genome using HISAT2. Bowtie2 was used to align clean reads to a comprehensive gene set, including both known and novel coding and noncoding transcripts. Gene expression levels were quantified using RSEM (v1.3.1). The differential gene expression and gene set enrichment analyses were performed in the R environment.

1.3 Lipids analysis and lipoprotein profile measurement

Mice were fasted overnight prior to blood collection. Plasma was separated by centrifugation and stored at -80°C. Cell samples were collected and lysed following the manufacturer’s instructions. Total cholesterol and triglycerides were determined using the Total Cholesterol Assay Kit and Triglyceride Assay Kit (Nanjing Jianjian Bioengineering Institute). Plasma HDL-cholesterol and LDL-cholesterol concentrations were determined using the HDL-cholesterol and LDL-cholesterol Assay Kits (Nanjing Jianjian Institute of Bioengineering, Nanjing, China) following the provided protocols. Palmitic acid levels in cells were measured using the Human Palmitic Acid ELISA Kit (MeiMian) according to the manufacturer’s instructions.

1.4 Western blotting analysis

Samples were lysed in RIPA buffer (Beyotime Biotechnology) containing protease and phosphatase inhibitors. Protein concentration was determined using the BCA Protein Assay Kit (Beyotime Biotechnology). Equal amounts of protein were separated by SDS-PAGE and transferred to a polyvinylidene difluoride (PVDF) membrane. The membrane was blocked with 5% skimmed milk in Tris-buffered saline containing 0.1% Tween-20 (TBST) and incubated overnight at 4°C with the primary antibody at the recommended dilution (detailed in Table S2). After three 10-minute washes with TBST, the membrane was incubated with horseradish peroxidase-conjugated secondary antibody for 1 hour at room temperature. Chemiluminescent signals were captured using the Amersham Imager 600 (GE Healthcare), and band intensity was analyzed using ImageJ.

1.5 RT-qPCR analysis

Total RNA was isolated from cells or tissues using AG RNAex Pro reagent (Accurate Biotechnology). One microgram of RNA was reverse transcribed to cDNA using the Evo M-MLVRT kit (Accurate Biotechnology). Quantitative PCR was performed using the LightCycler 480 II thermal cycler (Roche) with SYBR Green Premix Pro Taq HS (Accurate Biotechnology). Relative gene expression levels were normalized to β-actin and calculated using the 2-ΔΔCt method. Primer sequences are listed in Supplemental Table S3.

| 2. Supplementary tables  Table S1 Baseline characteristics of patients (n=12) | | | | | | |
| --- | --- | --- | --- | --- | --- | --- |
|  | gender | age | smoking | hypertension | T2DM | CAD |
| donor 1 | male | 53 | no | no | no | yes |
| donor 2 | male | 49 | no | yes | no | no |
| donor 3 | female | 45 | no | no | no | no |
| donor 4 | female | 38 | no | no | no | no |
| donor 5 | male | 32 | no | no | no | no |
| donor 6 | male | 40 | no | no | no | no |
| patient 1 | male | 52 | yes | no | yes | no |
| patient 2 | male | 59 | no | yes | yes | yes |
| patient 3 | male | 65 | yes | yes | no | no |
| patient 4 | female | 93 | no | yes | yes | yes |
| patient 5 | male | 75 | yes | yes | no | yes |
| patient 6 | male | 68 | yes | yes | no | yes |

T2DM: type 2 diabetes mellitus; CAD: coronary artery disease

| Table S2 List of antibodies |  |  |
| --- | --- | --- |
| Antibodies | Source | Catalogue number |
| Rabbit anti-TMEM41B (1:1000 for WB) | Proteintech | 29270-1-AP |
| Rabbit anti-TMEM41B (1:100 for IF) | Invitrogen | PA5-143837 |
| Mouse anti-FASN (1:1000 for WB, 1:200 for IF) | Proteintech | 66591-1-Ig |
| Rabbit anti-OCT-1 (1:50 for CHIP) | Cst | 8157 |
| Rabbit anti-CD36 (1:1000 for WB) | Proteintech | 18836-1-AP |
| Rabbit anti-CD204 (1:1000 for WB) | Abcam | ab314227 |
| Rabbit anti-LOX1 (1:1000 for WB) | Proteintech | 11837-1-AP |
| Rabbit anti-ABCA1(1:1000 for WB) | Proteintech | 26564-1-AP |
| Mouse anti-Beta Actin (1:5000 for WB) | Proteintech | 66009-1-Ig |
| Rabbit anti-DDDDK tag (1:1000 for WB, 1:30 for IP) | Abcam | ab205606 |
| Rabbit anti-HA-Tag (1:1000 for WB, 1:50 for IP) | Cst | 3724 |
| Mouse anti-alpha smooth muscle actin (1:200 for IF) | Abcam | ab7817 |
| Rabbit F4/80 (1:200 for IF) | Proteintech | 28463-1-AP |
| Mouse anti-His-Tag (1:5000 for WB) | Proteintech | 66005-1-Ig |
| Alexa 488 Goat anti-Rabbit (1:1000) | Thermo Fisher | Cat# A-11008 |
| Alexa 568 Goat anti-Mouse (1:1000) | Thermo Fisher | Cat# A-11031 |
| HRP-conjugated Goat Anti-Rabbit IgG(H+L) (1:10000) | Proteintech | SA00001-2 |
| HRP-conjugated Goat Anti-Mouse IgG(H+L) (1:10000) | Proteintech | SA00001-1 |

| Table S3 Primer sequence for RT-qPCR | | | | |
| --- | --- | --- | --- | --- |
| **name** | |  | | **sequences** |
| Primers for IL1β | Forward (5’ - 3’) | | ATGATGGCTTATTACAGTGGCAA | |
|  | Reverse (5’ - 3’) | | GTCGGAGATTCGTAGCTGGA | |
| Primers for IL6 | Forward (5’ - 3’) | | ACTCACCTCTTCAGAACGAATTG | |
|  | Reverse (5’ - 3’) | | CCATCTTTGGAAGGTTCAGGTTG | |
| Primers for TNFα | Forward (5’ - 3’) | | CCTCTCTCTAATCAGCCCTCTG | |
|  | Reverse (5’ - 3’) | | GAGGACCTGGGAGTAGATGAG | |
| Primers for TMEM41B | Forward (5’ - 3’) | | TTGTGTTCTGGACTTGGTGC | |
|  | Reverse (5’ - 3’) | | TCTATGACGTTCAACCTGCTGT | |
| Primers for β-Actin | Forward (5’ - 3’) | | TTTCTGTCACTCTTCTCTTAGGT | |
|  | Reverse (5’ - 3’) | | AGGTCTTTACGGATGTCAAGG | |

| Table S4 The primers specific for the TMEM41B promoter | | | | |
| --- | --- | --- | --- | --- |
| **name** |  | | | **sequences** |
| Primers for site NC inTMEM41B promoter | | Forward (5’ - 3’) | GAAACTGATGATTGGCAGCT | |
|  | | Reverse (5’ - 3’) | TCTCTTATGTCTACTTCTTTCTAC | |
| Primers for site 1 inTMEM41B promoter | | Forward (5’ - 3’) | GAGTTGTGCTTTCCTTACTG | |
|  | | Reverse (5’ - 3’) | TATATTTATGGGCCAGGTGC | |
| Primers for site 2 inTMEM41B promoter | | Forward (5’ - 3’) | GGATGTGTACATAAAACAGT | |
|  | | Reverse (5’ - 3’) | CTATTCAGTAATTACATTTG | |
| Primers for site 3 inTMEM41B promoter | | Forward (5’ - 3’) | AAGCTAAATGTAGAGCTGAC | |
|  | | Reverse (5’ - 3’) | CTAAGGCTCAGAGAGGGCTA | |

| Table S5 TMEM41B siRNA sequences | | |
| --- | --- | --- |
| **name** |  | **sequences** |
| si-TMEM41B#1(mouse) | Forward (5’ - 3’) | GAAUAUGAAGGUUCCGAGA(dT)(dT) |
|  | Reverse (5’ - 3’) | UCUCGGAACCUUCAUAUUC(dT)(dT) |
| si-TMEM41B#2(mouse) | Forward (5’ - 3’) | GCAAGAACAUCACUCCUUA(dT)(dT) |
|  | Reverse (5’ - 3’) | UAAGGAGUGAUGUUCUUGC(dT)(dT) |
| si-TMEM41B#3(mouse) | Forward (5’ - 3’) | CGUCACAGAGAACAUCUUA(dT)(dT) |
|  | Reverse (5’ - 3’) | UAAGAUGUUCUCUGUGACG(dT)(dT) |
| si-TMEM41B#1(human) | Forward (5’ - 3’) | GUUUCCUGGAACUCAAUAU(dT)(dT) |
|  | Reverse (5’ - 3’) | AUAUUGAGUUCCAGGAAAC(dT)(dT) |
| si-TMEM41B#2(human) | Forward (5’ - 3’) | GGUUGAACGUCAUAGAGAA(dT)(dT) |
|  | Reverse (5’ - 3’) | UUCUCUAUGACGUUCAACC(dT)(dT) |
| si-TMEM41B#3(human) | Forward (5’ - 3’) | GCAGGAACAACACUGUAUCAA(dT)(dT) |
|  | Reverse (5’ - 3’) | UUGAUACAGUGUUGUUCCUGC(dT)(dT) |

| Table S6 TMEM41B shRNA sequences | | | | |
| --- | --- | --- | --- | --- |
| **name** |  | | **sequences** | |
| shRNA-TMEM41B#1 | | sense (5’ - 3’) | | TTCAGATGATCTTTGCTGCCA |
|  | | antisense (5’ - 3’) | | TGGCAGCAGATCATCTGAA |
| shRNA-TMEM41B#2 | | sense (5’ - 3’) | | TAGGAGAGCATGTAGCAGAAT |
|  | | antisense (5’ - 3’) | | ATTCTGCTATGCTCTCCTA |
| shRNA-TMEM41B#3 | | sense (5’ - 3’) | | ATCAGAATAAATACCGAGCTC |
|  | | antisense (5’ - 3’) | | GAGCTCGGTTTATTCTGAT |

3. Supplementary figures

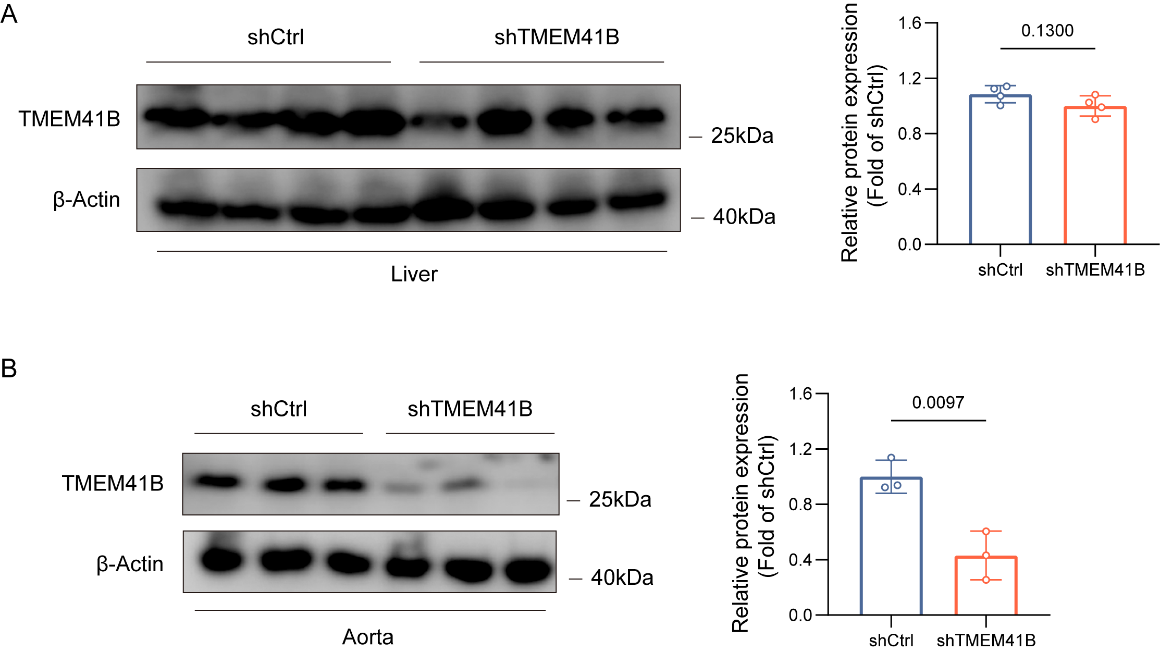

Fig. S1 Validation of TMEM41B knockdown in the aorta.

(A) Western blot analysis demonstrates no significant change in TMEM41B protein levels between shCtrl and shTMEM41B groups in the liver (n = 4 per group). (B) TMEM41B protein levels are significantly reduced in the aorta of shTMEM41B mice compared to shCtrl mice (n = 3 per group). Data are presented as mean ± SD and analyzed using an unpaired, two-tailed Student’s t-test.

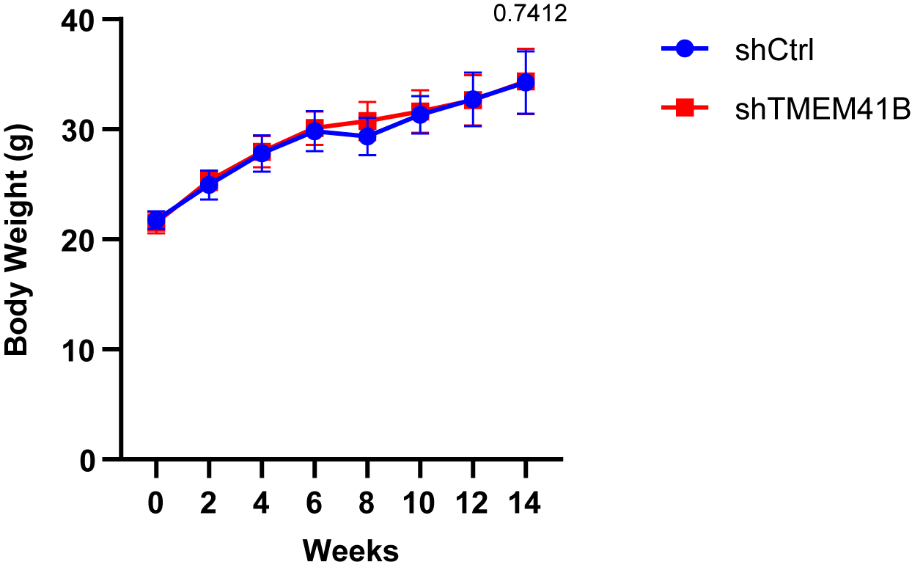

Fig. S2 Body weight changes in AAV-shCtrl and AAV-TMEM41B-injected mice.

Body weight was monitored in ApoE-/- mice injected with AAV-shCtrl or AAV-TMEM41B and fed a Western diet. Measurements were taken at regular intervals (n = 10 mice per group). Data are presented as mean ± SD and analyzed using two-way ANOVA.

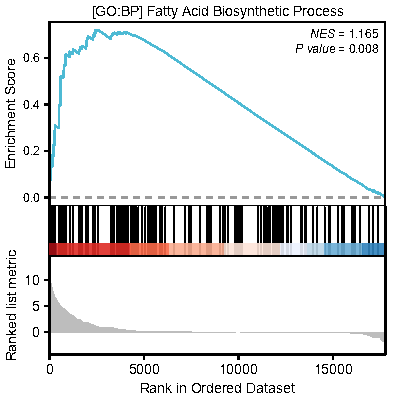

Fig. S3 TMEM41B modulates lipid profiles in VSMCs.

Gene Set Enrichment Analysis (GSEA) reveals a global upregulation of the fatty acid biosynthetic process in oe-TMEM41B compared to oe-Ctrl in VSMCs.

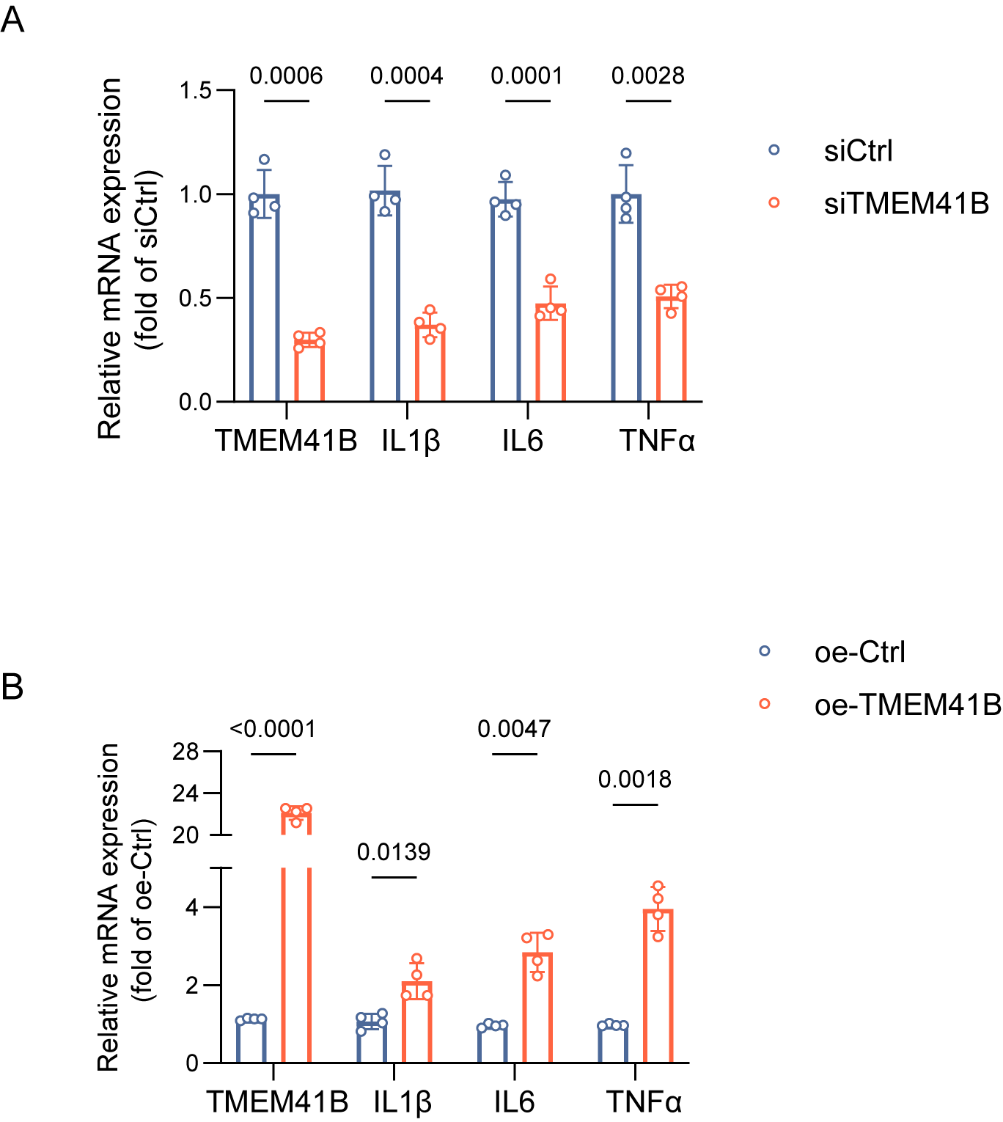

Fig. S4 TMEM41B increases the production of inflammatory factors.

(A) RT-qPCR analysis of IL1β, IL6, and TNFα mRNA expression in HASMCs transfected with siCtrl or siTMEM41B (n = 4). (B) RT-qPCR analysis of IL1β, IL6, and TNFα mRNA expression in HASMCs transfected with siCtrl or siTMEM41B (n = 4). All data are presented as mean ± SD and analyzed using an unpaired, two-tailed Student’s t-test.

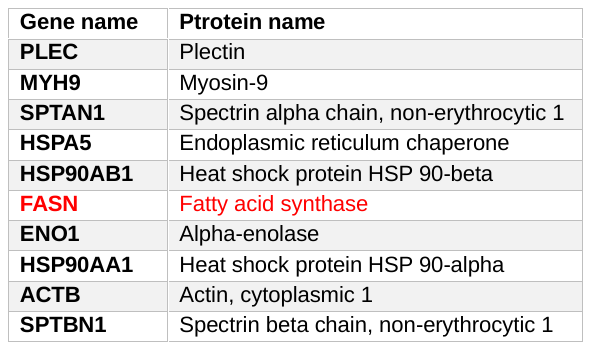

Fig. S5 Top 10 protein candidates identified by IP-MS.

HASMCs were transfected with HA-TMEM41B, and 48 hours post-transfection, immunoprecipitation (IP) and mass spectrometry (MS) were performed to identify interacting proteins.

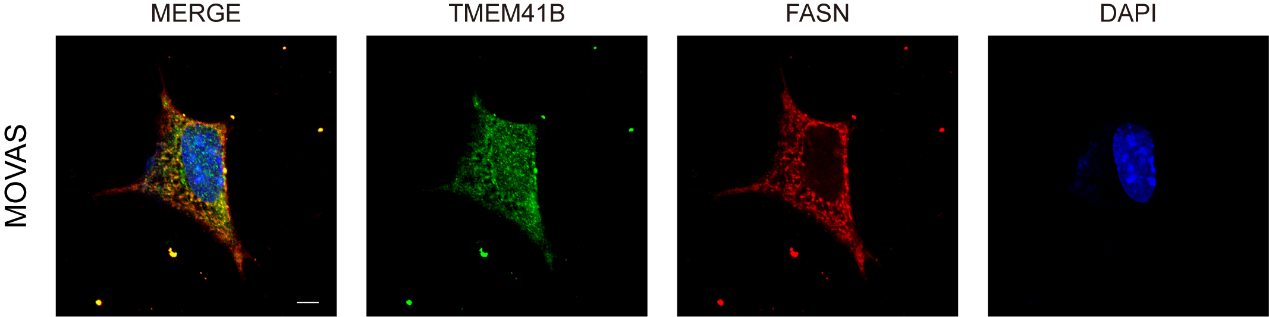

Fig. S6 Colocalization of TMEM41B and FASN in MOVAS cells.

MOVAS cells were immunofluorescently stained with anti-TMEM41B (green) and anti-FASN (red) antibodies. Nuclei were counterstained with DAPI. Scale bar: 10 µm.

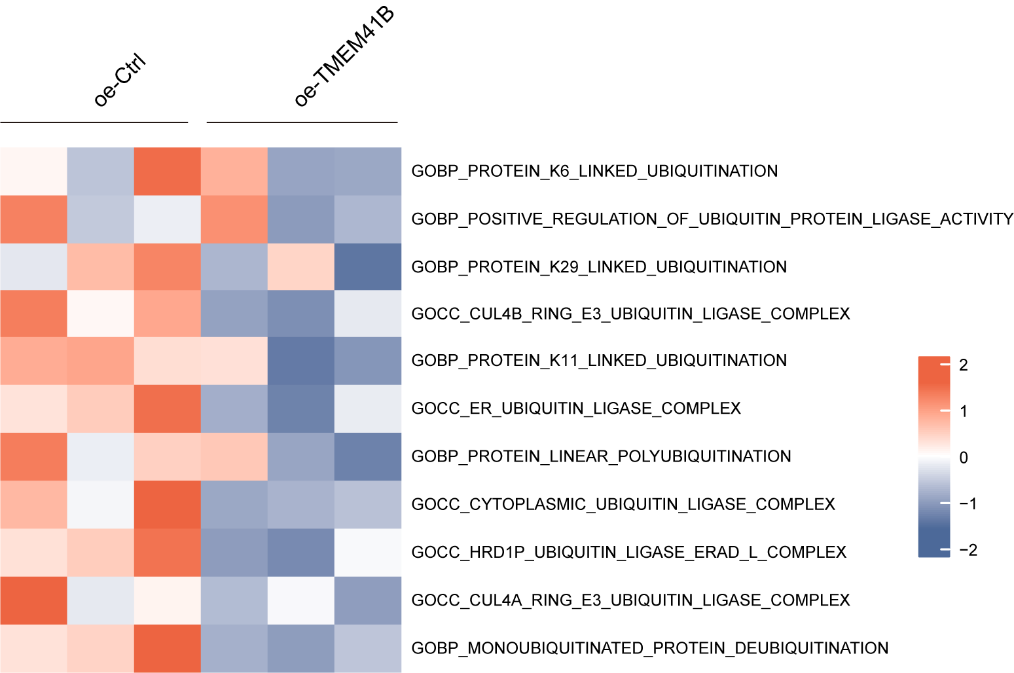

Fig. S7 Ubiquitination is downregulated in TMEM41B overexpression

GSVA analysis revealed a significant upregulation of ubiquitination-related gene sets in the oe-TMEM41B group
